# Interaction hierarchy among Cdv proteins drives recruitment to membrane necks

**DOI:** 10.1101/2024.10.03.616404

**Authors:** Nicola De Franceschi, Alberto Blanch-Jover, Cees Dekker

## Abstract

Cell division in the crenarchaea is accomplished by the Cdv system. In Sulfolobus cells, it was observed that an initial non-contractile ring of CdvA and CdvB forms at the mid location of the cell, which is followed by a second ring of CdvB1 and CdvB2 that appear to drive the constriction of the cell membrane. Here, we use an *in vitro* reconstituted system to explore how protein interactions among these Cdv proteins govern their recruitment to the membrane. We show that CdvA can bind to lipid membranes, but does so more efficiently when it is in complex with CdvB. We find that CdvB2 can polymerize if its self-inhibitory domain is removed, and that by itself is exhibits poor binding to the membrane. However, CdvB2 can be efficiently recruited to the membrane by both CdvB1 and CdvB. Furthermore, the CdvB1:CdvB2 co-polymer can be recruited to the membrane by CdvA:CdvB. By reconstituting these proteins in dumbbell-shaped liposomes, we show that Cdv proteins have a strong preference to localize at membrane necks of high curvature. Our findings clarify many of the mutual protein interactions of the Cdv system and their interaction with the membrane, thus helping to build a mechanistic understanding of cell division in archaeal cells.

## Introduction

Cell division in the archaeal phylum of the crenarchaea is performed by the Cdv system (Lindås et al., 2008; Samson et al., 2008). While this protein machinery is unique to archaea, some of its components are homologous to the ESCRT-III proteins that are responsible for the cell division, vesicle budding, and many other reverse-topology membrane-scission processes in eukaryotes (Schöneberg et al., 2016; Hurley, 2015). This homology is one of the many similarities that eukaryotes share with archaea, which reinforces the widespread idea that these two kingdoms of life share an evolutionary root (Cox et al., 2008).

Since the Cdv system was first described 15 years ago in Sulfolobus sulfataricus, it has been shown to be present in many other species of the TACK superphylum of archaea (Makarova et al., 2010). It is composed of CdvA, the CdvB paralogs (homologous to ESCRT-III in eukaryotes), and CdvC (homologous to the eukaryotic Vps4) (Samson et al., 2008). The first cell imaging of the Cdv system showed a ring of CdvA and CdvB forming at the center of the cell between two segregated nucleoids during division, which over time colocalized with a ring of CdvC in the same position (Lindås et al., 2008). All these 3 proteins are located in the same operon (Lindås et al., 2008), while paralogs of CdvB (namely CdvB1, CdvB2 and CdvB3) are located in other parts of the genome. Recently, it was reported that these paralogs play a crucial role in cell division (Tarrason Risa et al., 2020). A recent model for cell division in crenarchaea is that CdvA and CdvB initially form a ring at the centre of the cell, where it subsequently recruits CdvB1 and CdvB2 (Tarrason Risa et al., 2020). At this point, CdvB is digested by the proteasome and the initial CdvA:CdvB ring gets removed from the membrane, while CdvB1 and CdvB2 are left to perform the constriction of the membrane, presumably through the interaction of CdvB1 directly with the membrane (Blanch Jover et al., 2022), until the final step of scission (Tarrason Risa et al., 2020). CdvC is a AAA ATPase that has been suggested to remove monomers of CdvB1 and CdvB2 from the ring, generating a turnover that ensures cellular constriction while avoiding steric hindrance at the neck (Tarrason Risa et al., 2020; Harker-Kirschneck et al., 2022). This model of action of the Cdv proteins is supported by some experimental findings on live cells. When generating mutant cells of S. sulfataricus lacking CdvB1, these presented a normal constriction of the membrane, but some cells failed to perform the last step of scission, leaving them with two full copies of the genome (Pulschen et al., 2020). Furthermore, cells lacking CdvB2 were able to perform the scission normally, but tended to present a misplacement of the constricting ring, resulting in aberrant daughter cells that were not equally sized (Pulschen et al., 2020). This indicates that the two paralogs responsible for the constriction actually perform different roles during the division process. However, it also shows how cells with any of these two proteins can still perform, albeit with difficulties, the full process of constriction and scission on their own.

While a global picture has thus been emerging, many of the underlying mechanistic interactions remain unclear. CdvA has an E3B domain (ESCRT-III binding region; see Figure 1A) through which it is capable of interacting with the wH (winged helix) region of CdvB (Moriscot et al., 2011). None of the other paralogs present such a wH domain (Caspi et al., 2018). Additionally, CdvA can bind to lipid membranes while CdvB is not able to do so (Samson et al., 2011). Therefore, CdvA is seen as the membrane recruiter of CdvB to the membrane. In turn, CdvB is known to interact with CdvB1, which then interacts with CdvB2 (Samson et al., 2008), suggesting that CdvB is the recruiter of the CdvB1:CdvB2 constricting ring. Finally, during the constriction of the membrane, CdvC is presumed to disassemble the filaments of the CdvB1:CdvB2 polymer to generate a turnover of protein and thus supply energy to the system to deform the membrane (Harker-kirschneck et al., 2022). This ATPase features major structural similarities to the eukaryotic Vps4 (Caillat et al., 2015), which is known to be responsible for the depolymerization of ESCRT-III filaments (Lata et al., 2008; Azad et al., 2023) and to create a turnover of ESCRT-III components at the membrane (Chiaruttini et al., 2015; Pfitzner t al., 2020). In archaea, this similarity is further strengthened by in vitro experiments where CdvC was able to depolymerize CdvB1 filaments and detach CdvB1 from lipid membranes into solution (Blanch Jover et al., 2022).

**Figure 1:**
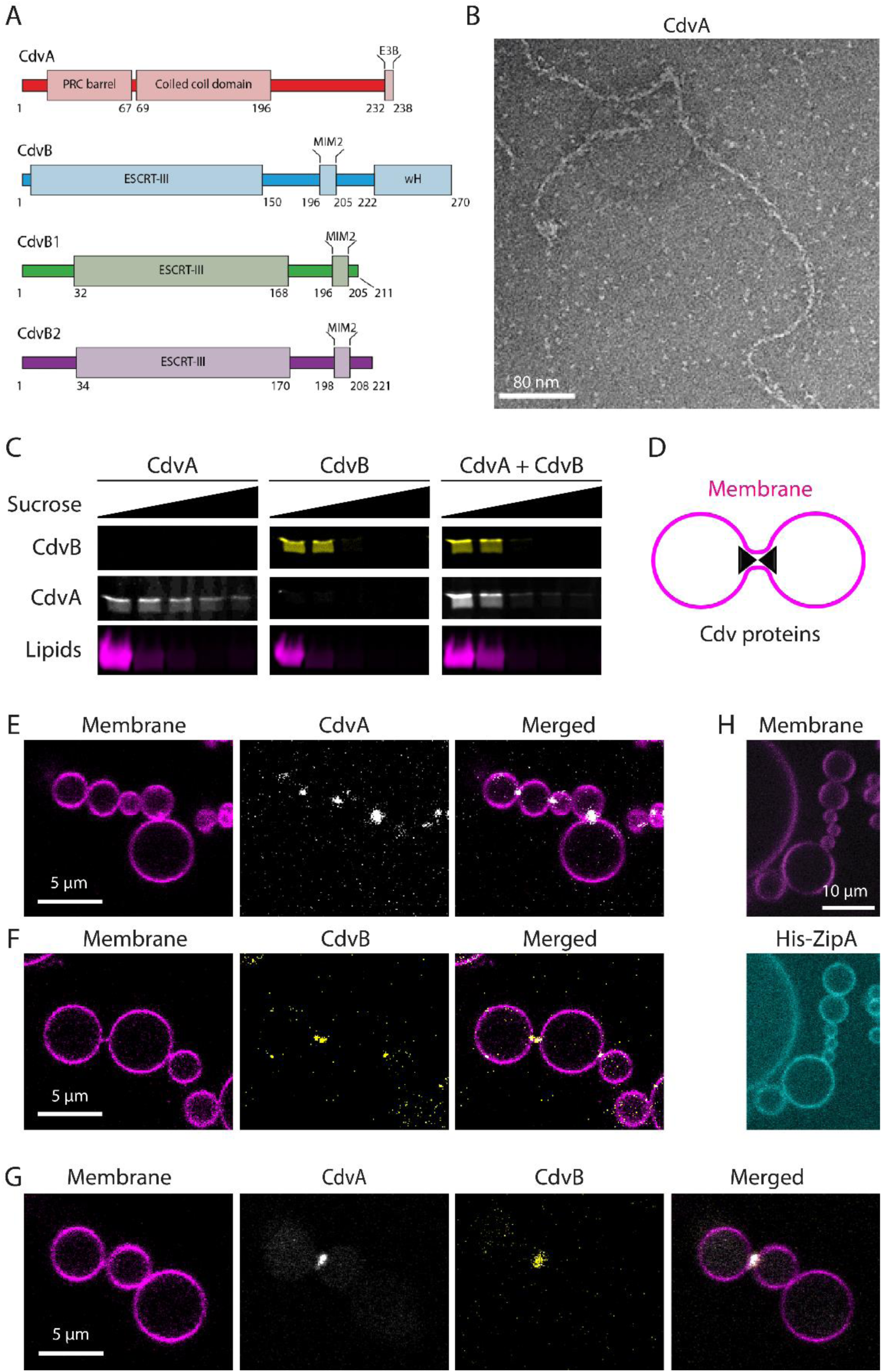
polymerization and membrane binding of CdvA and CdvB. (A) Schematic depicting the domain architecture of Cdv proteins. (B) Negative staining TEM image of elongated filaments formed by CdvA upon removal of the MBP tag in the absence of DNA. (C) Flotation assay showing recombinant Cdv proteins binding to multilamellar vesicles. CdvA alone is distributed in all fractions, while CdvB alone is found predominantly in the liposome-bound fraction. When both CdvA and CdvB are present, both proteins are found predominantly in the liposome-bound fraction. A representative gel from 3 independent experiments is shown. (D) Schematic representing the topology of Cdv proteins reconstituted inside dumbbell-shaped liposomes, recapitulating the shape of a dividing cell. (E) Confocal image of CdvA reconstituted inside dumbbell-shaped liposomes. CdvA localizes at the neck. A representative image from 2 independent experimental preparations is shown. (F) Representative confocal image of CdvB reconstituted inside dumbbell-shaped liposomes. CdvB localizes at the neck. A representative image from 2 independent experimental preparations is shown.(G) Representative confocal image of the CdvA:CdvB complex reconstituted inside dumbbell-shaped liposomes. Both proteins localize at the neck. A representative image from 2 independent experimental preparations is shown. (H) Binding of His-tagged ZipA to dumbbell liposomes containing 18:1 DGS-NTA lipids. As expected for a protein binding the membrane via a His-tag, ZipA exhibits a homogeneous membrane distribution, without enrichment at the neck.

Little is yet known about how the Cdv proteins are hierarchically recruited to the membrane and how their mutual interactions affect this process. Here, we explore these questions using a bottom-up reconstitution approach. We find that CdvA can bind to lipid membranes, but does so more efficiently when it is interacting with CdvB. We observe that CdvB2 can be recruited to the membrane by both CdvB1 and CdvB. Finally, we show that all Cdv proteins exhibit a preferential binding for highly curved membranes and preferentially localize at the necks of dumbbell-shaped vesicles.

## Results

### Filament formation by CdvA

Previous in vitro studies of purified CdvA were done with either a full-length version of the protein from M. sedula (Moriscot et al., 2011) or with an N-terminus-truncated CdvA from S. acidocaldarius that was missing the initial PRC barrel (Figure 1A) (Samson et al., 2011; Dobro et al., 2013). The phenotype of these two versions differed in that the full-length protein formed double helical filaments that were reported to be stabilized by the binding of DNA, whereas the N-terminus-truncated CdvA did not polymerize. At the same time, the N-terminus-truncated CdvA was shown to be able to bind to lipid membranes and recruit CdvB to the membrane along with it. For our study, we purified full length CdvA from M. sedula, following the protocol published by Moriscot et al., 2011, and we obtained the same type of filaments as described in their work (Supplementary Figure 1A). However, we never observed interaction between CdvA and lipid membranes (data not shown). We then decided to fuse the full length CdvA to an MBP-tag. The resulting purified MBP-CdvA formed short and thick polymers (Supplementary Figure 1B), which had an average length of 90 ± 40 nm (mean ± SD, N=195) and a width of 12 ± 3 nm (mean ± SD, N=156). When treating the protein with a TEV protease that cleaved the MBP from the protein, CdvA polymerized into long and thin filaments (Figure 1B) with an average length of 220 ± 100 nm (mean ± SD, N=102) and a width of 5 ± 1 nm (mean ± SD, N=106). Formation of these filaments did not require addition of DNA. Thus, the presence of the MBP tag allowed to control the polymerization of CdvA.

### CdvB enhances CdvA binding to the membrane

Next, we sought to investigate the lipid-binding capabilities of the full-length CdvA and CdvB. For this, we used a liposome flotation assay, where protein was mixed with negatively charged multilamellar vesicles (see Methods), added to a sucrose gradient, and spun at high speed. This yielded multiple fractions: upper fraction(s) containing liposomes (1), intermediate fraction(s) containing the soluble unbound protein, and the bottom fraction containing protein filaments sedimented at the bottom of the tube. In these assays, we mixed MBP-CdvA with the lipids together with TEV protease, in order to cleave the MBP tag. When testing for the binding of CdvA to lipid membrane, we observed that the protein was distributed to all fractions, including the lipid-bound fraction (Figure 1C). We then purified CdvB which, as previously shown (Moriscot et al., 2011), presented no filamentation regardless of having the MBP-tag or not. In liposome flotation assay, CdvB was highly enriched in the liposome-bound fraction (Figure 1C). In contrast, CdvB from S. acidocaldarius (called Saci_1373) was reported to not show membrane binding on its own (Samson et al., 2011). Thus it appears that CdvB from M. sedula may differ in this regard, although this difference could also be ascribed to different experimental conditions. When CdvA was mixed with CdvB, both proteins were found primarily in the liposome-bound fraction (Figure 1C). This indicates that the binding of full-length CdvA to the lipid membrane is enhanced by CdvB.

The human ESCRT-III system preferentially locates at membrane necks that present high curvatures (De Franceschi et al., 2018; Bertin et al., 2020; Pfitzner et al., 2020). We explored if a similar preference for high curvatures could be observed for Cdv proteins as well. Here, we used a recently developed SMS approach to produce dumbbell-shaped liposomes (De Franceschi et al., 2022). Briefly, a lipid mixture including negatively charged lipids in chloroform was dried out and resuspended in oil. In this oil, we formed water-in-oil droplets in which the proteins were encapsulated, and these droplets were then left to sedimented by gravity through a lipid interface into an outer water phase, thus obtaining unilamellar liposomes that contained the protein on the inside. The outer phase was made of buffer containing DNA “nanostars” that anchor to the membrane, inducing curvature thereby generating dumbbells. We thus obtain chains of dumbbell-shaped liposomes that are mutually connected through membrane necks. Protein that are encapsulated within these liposomes can be studied for their binding in a membrane geometry that resembles cells that are dividing (Figure 1D). Such a system was recently used to demonstrate a membrane scission activity of Dynamin A (De Franceschi et al., 2024).

In these dumbbell-shaped liposomes, we observed a clear preference of both CdvA, CdvB and the CdvA:CdvB complex to localize at the membrane necks (Figure 1E, 1F, 1G). Proteins displayed a strong fluorescence signal at the membrane necks, while they showed only a residual weak homogeneous binding to other membrane regions of the dumbbell-shaped liposomes. We confirmed that the lobes of these dumbbell-shaped liposomes are connected by membrane necks (Movie 1). We also performed control experiments with a protein that bound the membrane, containing lipids functionalized with Ni-NTA, via a His-tag (ZipA from E. coli) and that was not expected to sense curvature. As expected, ZipA did not show any significant enrichment of the protein signal at the necks (Figure 1H and Supplementary Figure 1C), indicating that the enrichment on highly curved membranes is a specific property of the Cdv proteins and not an artifact induced by the SMS assay.

### Filaments formation by CdvB1 and CdvB2ΔC

Next, we purified CdvB2 from M. sedula fused to an MBP-tag. The resulting protein was not presenting any spontaneous filamentation either with or without the MBP tag (Supplementary Figure 2A, 2B). Moreover, we found that full-length CdvB2 was unable to bind membranes in liposome binding assays, neither alone nor in combination with CdvB1 (Supplementary Figure 2C). ESCRT-III proteins commonly feature an inactive soluble state and an active membrane-bound state (Tang et al., 2015). Indeed, *in vitro*, these proteins remain inactive soluble monomers while they may get activated and able to polymerize by deleting their C-terminus part (Shim et al., 2007). The same has been previously shown for purified CdvB (Moriscot et al., 2011), which did not present any polymerization *in vitro*, but spontaneously assembled into filaments upon removal of its C-terminus. Hence, we decided to explore if the same was true for CdvB2, and we made a C-terminal truncated version that contained amino acids 1-170 of CdvB2, henceforth denoted as CdvB2ΔC.

TEM imaging of the CdvB2ΔC-MBP fusion protein showed that it was polymerizing into a characteristic shape of well-defined short linear filaments (Supplementary Figure 2D). Similarly to CdvA, CdvB2ΔC assembled into longer and thinner filaments upon MBP tag cleavage (Figure 2A and Supplementary Figure 2E, 2F), of average length 245 ± 95 nm (mean ± SD; N=36) and width of 14 ± 3 nm (mean ± SD; N=64). These data indicate that CdvB2 also presents a self-inhibitory domain, and that filament formation can be triggered by its removal.

**Figure 2:**
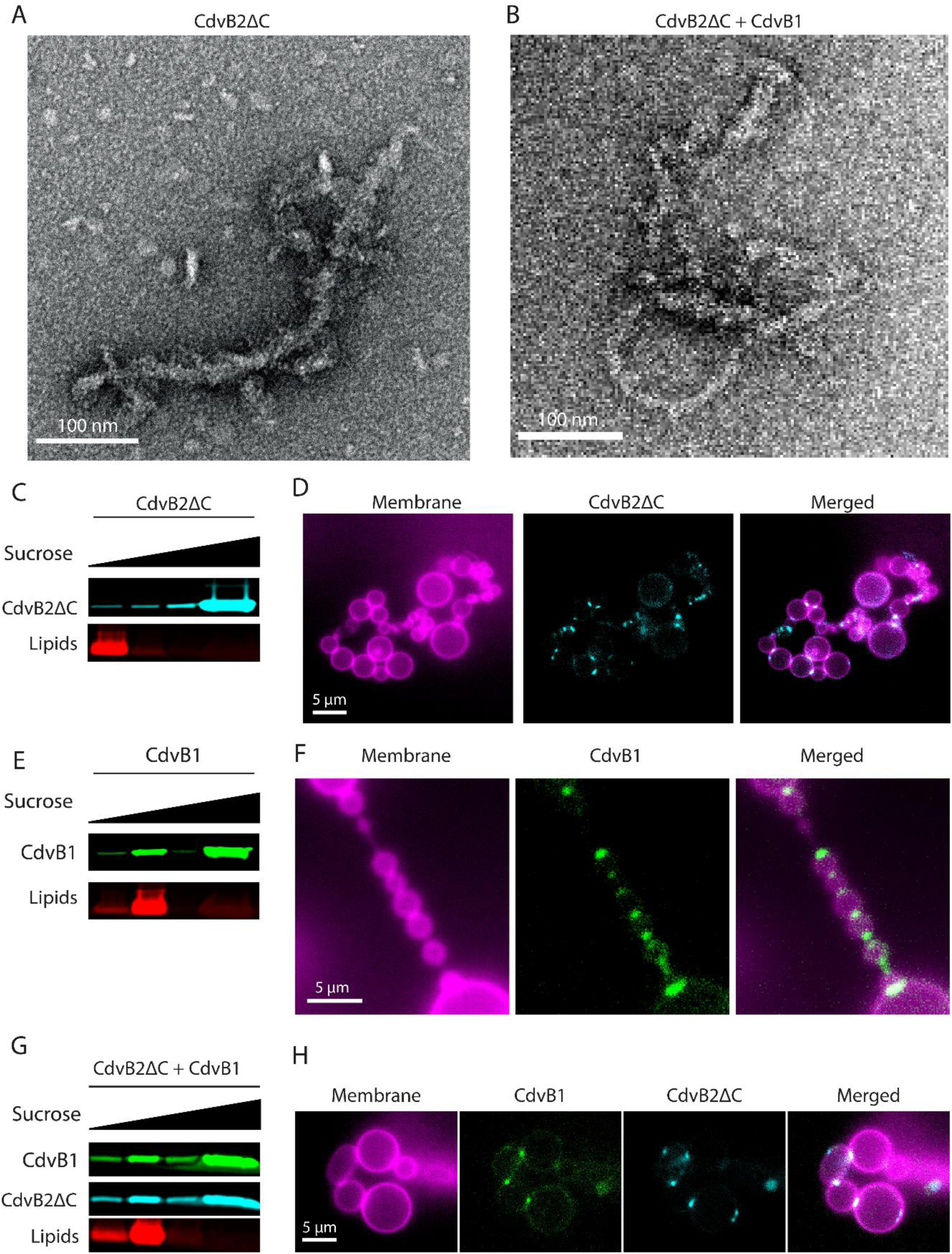
polymerization and membrane binding of the CDVB1: CdvB2ΔC complex. (A) Negative staining TEM image of CdvB2ΔC filaments obtained upon removal of the MBP tag. (B) Negative staining TEM image of CdvB1:CdvB2ΔC co-polymer. (C) Liposome flotation assays showing very limited membrane binding of CdvB2ΔC alone. A representative gel from 2 independent experimental preparations is shown. (D) Microscopy image of fluorescently labelled CdvB2ΔC reconstituted inside dumbbell-shaped liposomes and localizing at necks. (E) Liposome flotation assays showing clear membrane binding of CdvB1 alone. A representative gel from 2 independent experimental preparations is shown. (F) Microscopy image of fluorescently labelled CdvB1 reconstituted inside dumbbell-shaped liposomes and localizing at necks. (G) Liposome flotation assays showing CdvB2ΔC being recruited to membranes along with CdvB1. A representative gel from 2 independent experimental preparations is shown. (H) Microscopy image of a fluorescently labelled CdvB1:CdvB2ΔC complex reconstituted inside dumbbell-shaped liposomes and localizing at necks.

Since CdvB2 forms part of the constricting ring together with CdvB1, we explored the effects of their interaction on filament formation. When mixing CdvB1 and CdvB2ΔC together, filaments formed with a very similar length (272 ± 94 (mean ± SD; N=84)) and width (15 ± 6 (mean ± SD; N=184)) to that of CdvB2ΔC alone (Figure 2B and Supplementary Figure 2E, 2F). Interestingly, these filaments appeared to be more curved. To quantify this, we measured the ratio of the end-to-end distance to the contour length of filaments, which equals 1 for perfectly straight filaments but is <1 for curved ones. For CdvB2ΔC filaments we measured a ratio of 0.86 ± 0.13, while the CdvB1:CdvB2ΔC co-polymer yielded a ratio of 0.75 ± 0.21. This may also imply that the CdvB1:CdvB2ΔC copolymer is more flexible than the CdvB2ΔC homopolymer.

### CdvB1 and CdvB2ΔC binding to membrane necks

Subsequently, we tested the ability of CdvB2ΔC to bind lipid membranes. When mixed with negatively charged multilamellar vesicles, CdvB2ΔC was almost exclusively found in the soluble and filamented fractions (Figure 2C), with only a small fraction of the protein found in the lipid-bound fraction. In spite of its low affinity, we encapsulated CdvB2ΔC in dumbbells using the SMS technique and observed instances of CdvB2ΔC clusters at their neck (Figure 2D). It is possible that, in the presence of the correct membrane curvature found at the neck of dumbbells, the affinity of CdvB2ΔC for membrane may increase, indicating that CdvB2ΔC is able to sense curvature. CdvB1 presented clear membrane binding when mixed with liposomes on its own (Figure 2E), as we previously showed (Blanch Jover et al., 2022). CdvB1 also exhibited preferential neck localization when reconstituted in dumbbells (Figure 2F). When mixing the two proteins and subsequently adding them to multilamellar vesicles, CdvB1 appears to recruit CdvB2ΔC to the membrane (Figure 2G), indicating that they form a complex. Accordingly, we observe binding of the CdvB1:CdvB2ΔC complex to the neck of dumbbell liposomes (Figure 2H).

### Reconstitution of a quaternary CdvA:CdvB:CdvB1:CdvB2ΔC complex at a membrane neck

Finally, we explored the interaction between the CdvA:CdvB and CdvB1:CdvB2ΔC complexes in the presence of lipid membranes. In flotation assays, CdvB1 was recruited to the membrane in the presence of CdvA and CdvB (Figure 3A and Supplementary Figure 3A). Moreover, the CdvA:CdvB complex was also able to recruit CdvB2ΔC to liposomes (Figure 3A). Finally, when mixing all four proteins together in the presence of liposomes, we observed that they were all recruited to liposomes (Figure 3A and Supplementary Figure 3B). Thus, our data indicate that either CdvB1 or a CdvA:CdvB complex (or both) can recruit CdvB2 to the membrane.

**Figure 3:**
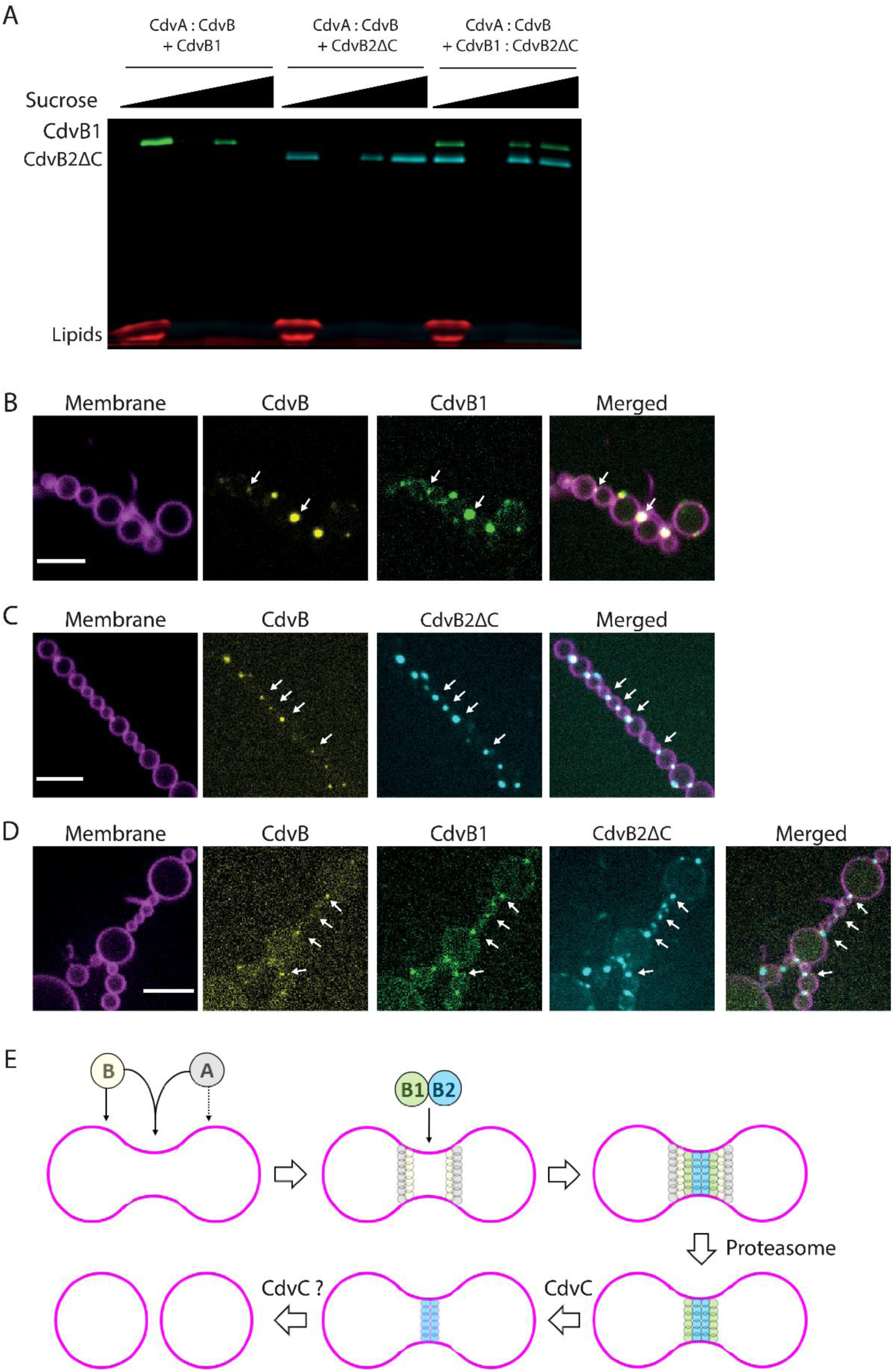
membrane binding of the CdvA:CdvB:CdvB1:CdvB2ΔC quaternary complex. (A) Liposome flotation assays showing recruitment of CdvB1 and CdvB2ΔC to membranes, either alone or in combination, by the CdvA:CdvB complex. (B) Spinning disk confocal images of the ternary complex CdvA (unlabelled) + CdvB-Alexa568 + CdvB1-Alexa488 reconstituted inside the neck of dumbbell liposomes. (C) Spinning disk confocal images of the ternary complex CdvA (unlabelled) + CdvB-Alexa568 + CdvB2ΔC-Cy5 reconstituted inside the neck of dumbbell liposomes. (D) Spinning disk confocal images of the quaternary complex CdvA (unlabelled) + CdvB-Alexa568 + CdvB1-Alexa488 + CdvB2ΔC-Cy5 reconstituted inside the neck of dumbbell liposomes. Scale bars: 10 µm. (E) Schematic depicting the stepwise assembly and disassembly of the Cdv division ring. CdvA and CdvB can bind the membrane individually, but binding of CdvA is enhanced by CdvB, and the two proteins are able to assemble at the neck by forming a complex. Once the CdvA:CdvB ring is assembled, CdvB1 and CdvB2 are both recruited at the neck. Subsequently, the proteasome removes CdvA, and CdvC removes CdvB1, leaving only CdvB2, which may be removed by CdvC, thus achieving membrane abscission.

We then reconstituted the same combinations of Cdv proteins inside dumbbells-shaped liposomes: CdvA (unlabelled) + CdvB + CdvB1 (Figure 3B); CdvA (unlabelled) + CdvB + CdvB2ΔC (Figure 3C) and CdvA (unlabelled) + CdvB + CdvB1 + CdvB2ΔC (Figure 3D). In passing we note that assembly of such a quaternary Cdv complex at membrane necks of dumbbell-shaped liposomes resembling the shape of a dividing cell has never been achieved before. In all cases, we observed a clear preference of the complexes to assemble onto regions of high membrane curvature, as demonstrated by the strong fluorescence signal at the membrane necks. In the quaternary complex, the fluorescence intensity of the Cdv proteins at the necks was about 7 times higher than that at the membrane away from the necks. For comparison, the lipid fluorescence intensity at the neck was only ∼2 times enhanced (Supplementary Figure 3C) as expected in the neck region where the membrane of the two lobes are in close proximity.

## Discussion

In this study, we elucidated the protein interaction network that governs Cdv protein assembly at the membrane, and showed that Cdv complexes spontaneously localize at membrane necks. We learned that full-length CdvA can form filaments without the need of DNA to stabilize them, and that it binds the membrane to some extent on its own. CdvB from M. sedula binds the membrane efficiently, unlike its homologous from S. acidocaldarius (Samson et al., 2011). When in complex with CdvB, CdvA binds the membrane more efficiently. This suggests that both proteins are likely recruited to the membrane jointly. We confirmed our previous finding that CdvB1 interacts with membranes on its own (Blanch Jover et al., 2022). Furthermore, we purified CdvB2 and showed that it is able to polymerize into filaments once the C-terminal domain is removed. This is a relevant parallelism with the ESCRT machinery, where many of the proteins also only polymerize upon removal of the C-terminus. We observed that CdvB2 binds poorly the membrane by itself, even upon removal of the C-terminal. However, we observed that either CdvB1 or a CdvA:CdvB complex (or both) can recruit CdvB2ΔC to the membrane, which suggests that CdvB2ΔC interaction with the membrane may require these proteins to be present. Intriguingly however, we also observe binding of CdvB2ΔC alone to highly curved membrane, which is the geometry found at the neck of dumbbells. Thus, it is also possible that the mechanism by which CdvB1 and the CdvA:CdvB complex recruit CdvB2ΔC at the membrane is by generating curvature. Further work is needed to thoroughly dissect this phenomenon for the various Cdv proteins.

Recently, in vivo imaging showed that CdvB1 is required to facilitate the spatial separation of CdvB and CdvB2 polymers, as absence of CdvB1 caused a spatial overlapping of CdvB and CdvB2 polymers at the intercellular bridge of dividing cells (Hurtig et al., 2023). Our *in vitro* data show that indeed CdvB2 can be recruited to the membrane by the CdvA:CdvB complex even in the absence of CdvB1. This suggests that the co-localization of CdvB and CdvB2 in the absence of CdvB1 *in vivo* is likely driven by direct interaction between CdvB and CdvB2.

Moreover, in simulations it was observed that, to obtain a non-overlapping arrangement of Cdv polymers, CdvB1 filaments should be relatively flexible compared to both CdvB and CdvB2 filaments (Hurtig et al., 2023). Our *in vitro* data are consistent with this scenario, as we observe that the CdvB1:CdvB2 co-polymers are more curved, and thus potentially more flexible, than the CdvB2 homo-polymers.

We found that all Cdv proteins preferentially localize at membrane necks, providing the first experimental evidence of affinity of Cdv proteins for high-curvature membranes. This presents another common trait between the archaeal and eukaryotic systems, as many ESCRT proteins also preferentially bind highly curved membranes (Lee et al., 2015; De Franceschi et al., 2018; Bertin et al., 2020; Pfitzner et al., 2020). Notably in our system, we cannot discriminate whether the proteins preferentially assemble on positive or negative curvature, because the neck present in dumbbells exhibits a catenoid geometry, which is characterized by both positive and negative curvature. Using super-resolution microscopy, it was shown that Cdv proteins arrange in ring-like structures that are not completely overlapping; that is, they take distinct spatial positions within the relatively highly curved membrane of the division ring (Hurtig et al., 2023). Our system is currently only amenable for standard confocal microscopy, due to constant moving of the dumbbells (De Franceschi et al., 2022). However, we can envision that protocols for fixation of dumbbells can be developed in the future, which would allow to investigate the assembly of recombinant proteins at the neck region using super-resolution microscopy. This would also help elucidating the nanostructure of CdvA at the neck. Our electron microscopy data show that CdvA assembles in rings that are below the diffraction limit, and accordingly we observe diffraction-limited clusters of CdvA and CdvB at necks. In contract, in vivo CdvA and CdvB were shown to form micrometer-sized rings (Hurtig et al., 2023). One possible interpretation for this discrepancy is that CdvA and CdvB would spontaneously prefer to assemble on a tighter neck, but in vivo they are prevented from doing so by additional factors. This provides a further parallelism with the eukaryotic ESCRT-II complex, in which some subunits may be forced to assemble in vivo in a suboptimal, “frustrated” geometry, and releasing this mechanical frustration could result in membrane deformation (Bertin, 2020).

Based on both literature and our own data, we can propose an updated model for the hierarchical assembly of Cdv proteins regulated by protein-protein interactions (Figure 3E): initially, CdvA and CdvB are recruited to the neck via their mutual interaction. Once the CdvA:CdvB complex forms a ring at the membrane, CdvB1 and CdvB2 are also recruited. In our experiment, CdvB1 and CdvB2 can bind the neck even in the absence of a pre-existing CdvA:CdvB complex at the membrane. However, the SMS system itself generates curvature (De Franceschi et al., 2022) and this may favour membrane binding, as our data suggest in the case of CdvB2. *In vivo*, recruitment of CdvB1 and CdvB2 may not occur unless a CdvA:CdvB complex is present. In all our experiments, we observe that CdvB1 and CdvB2ΔC always form a complex, whether in the presence of a membrane or not. Thus is it reasonable to assume that they are also recruited to the CdvA:CdvB complex together. Once the quaternary complex is assembled, CdvB is removed by the proteasome (Tarrason, 2020). In this regard, we provide the crucial piece of evidence that the CdvB1:CdvB2 complex can bind to membrane necks independently of the CdvA:CdvB complex. CdvB1 is subsequently removed from the membrane by CdvC (Blach-Jover, 2022), and here we show that CdvB2 can also bind to membrane necks by itself, without the presence of CdvB1.

We positioned the different subunits in our schematics (Figure 3E) based on their preferred radius of curvature proposed in (Hurtig et al., 2023), obtaining a final arrangement that resembles the later stage of constriction reported in (Hurtig et al., 2023). We expect this arrangement to be the case in our reconstituted system, even though the Cdv proteins are added simultaneously, because the shape of a neck is provided by the SMS system itself (De Franceschi et al., 2022) and therefore each protein can bind to the region having its preferred radius of curvature. This reconstituted system sets the stage to test Cdv-mediated membrane scission in the future, and in particular disassembly of CdvB1 and CdvB2 polymers from membrane necks by the action of CdvC. This will require the use of a sulfoscope (Pulschen et al., 2020), i.e., a microscope working at elevated temperatures, given that the catalytic activity of CdvC requires high temperature (Blanch Jover et al., 2022).

## Methods

### Plasmids

All of the proteins that we used are from Metallosphaera sedula., The original plasmids for CdvA (Msed_1670, UniProtID A4YHC3) and CdvB (Msed_1671, UniProtID A4YHC4) were kindly provided to us by Patricia Renesto’s lab. From those plasmids, the sequences of the proteins were copied and ordered as a synthetic gene already inserted in a pMAL-c5x from Biomatik, using BamHI and EcoRI cutting sites. Extra codons coding for cysteines were added at the N termini of the proteins for fluorescent labelling. The plasmid for CdvB1 was the same as used in our previous work (8). The gene for CdvB2 (Msed_1695, UniProtID A4YHE8) was obtained from the Gen Bank data base, and was reverse translated using the EMBOSS Backtranseq tool, optimized for E. coli codon usage. To the resulting DNA sequence, a codon of a cysteine for fluorescent labelling was added at the N terminal of the protein, as well as Tobacco Etch Virus (TEV) and an HRV 3C proteases cutting sites. The whole gene construct was ordered as a synthetic gene already inserted in a pMAL-c5x vector from Biomatik, using BamHI and EcoRI cutting sites.

From the original plasmid for MBP-CdvB2, the whole plasmid except for the C-terminus of the protein was copied by PCR. The resulting linearized plasmid was checked on an agarose gel, and then treated with the KLD reaction mix (New England Biolabs, Ipswich, Massachusetts, USA) to digest the template, phosphorylate the ends of the linearized plasmid and ligate it all at once. The reaction mix was then transformed into NEB5alpha competent cells (New England Biolabs), some colonies were picked, and the plasmid was purified using the QIAprep Spin Miniprep Kit (QIAgen, Hilden, Germany) and sent for sequencing.

#### 4.4.2. Protein purification

All proteins were produced in BL-21 E. coli strains. Cells were grown at 37°C in LBamp medium to an OD of around 0.5, at which point expression was induced with IPTG and cells were left to express the protein for 4 hours. After that, cells were harvested by centrifuging at 4500x g at 4°C for 12 minutes. For MBP-CdvA, the cell pellet was resuspended in lysis buffer (50mM Tris pH 8.8, 350 mM NaCl, 50 mM Glutamate, 50 mM Arginine, 0.05mM TCEP, cOmplete™ Protease Inhibitor Cocktail (Roche, Basel, Switzerland)), lysed by French press and centrifuged (150,000 g, 30 min, 4°C). The remaining supernatant was incubated with 1 ml of amylose resin (NEB, Ipswich, Massachusetts, USA) rotating for 2 hours at 4°C, after which it was poured through a gravity chromatography column and the protein was washed (50 mM Tris pH 8.8, 350 mM NaCl, 50 mM Glutamate, 50 mM Arginine, 0.05 mM TCEP) and eluted with elution buffer (50 mM Tris pH 8.8, 350 mM NaCl, 50 mM Glutamate, 50 mM Arginine, 0.05 mM TCEP, 10 mM maltose). For MBP-CdvB, MBP-CdvB2 and MBP-CdvB2ΔC, the cell pellet was resuspended in lysis buffer (50mM Tris pH 8.8, 50 mM NaCl, 0.05 mM TCEP, cOmplete™ Protease Inhibitor Cocktail) lysed by French press and centrifuged (150,000 g, 30 min, 4°C). The remaining supernatant was incubated with 1ml of amylose resin rotating for 2 hours at 4°C, after which it was poured through a gravity chromatography column and the protein was washed (50 mM Tris pH 8.8, 50 mM NaCl, 0.0 5mM TCEP) eluted with elution buffer (50 mM Tris pH 8.8, 50 mM NaCl, 0.05 mM TCEP, 10 mM maltose). MBP-CdvB1 was purified just as described in (8). After affinity chromatography, all proteins were run through a Superdex™ 75 increase 10/300 GL size exclusion chromatography column mounted in an ÄKTA™ Pure system. Samples were run with the same buffer as they were washed and stored by snap freeze in liquid nitrogen. Purity of the samples was evaluated by SDS PAGE stained with Coomassie blue. A fraction of all of the proteins was separated after the affinity chromatography and dialyzed into the same buffer but with pH of 7.4 to perform a maleimide-cysteine conjugation reaction. MBP-CdvA was labelled with Cy5, MBP-CdvB with Alexa-568, MBP-CdvB2ΔC with Cy5 and CdvB1 with Alexa 488. The rest of the purification stayed the same, and excess label was removed from the protein through the gel filtration column.

#### 4.4.3. TEM imaging

For imaging of MBP-CdvA, the protein was diluted down to 100 nM in buffer containing 50 mM Tris pH 7.4 and 50 mM NaCl (all samples were prepared using this buffer). For imaging the protein without MBP, 1μM of MBP-CdvA was mixed with 0.1 μM of TEV protease and left incubating at RT for 1 hour. The sample was then diluted 10 times before depositing it onto a carbon grid. MBP-CdvB2ΔC samples were diluted down to 100 nM in buffer, and samples without MBP were prepared by mixing 1 μM of MBP-CdvB2ΔC with 0.1 μM of TEV protease and left incubating at RT for 1 hour. Samples with CdvB1 and CdvB2ΔC were prepared by mixing MBP-CdvB1 and MBP-CdvB2ΔC both at 1μM concentration with 0.1 μM of TEV protease at RT for 1 hour. The samples were then diluted 10 times before depositing it onto a carbon grid. Samples were absorbed on glow-discharged carbon-coated 400-meshh copper grid purchased from Quantifoil (Großlöbichau, Germany) and stained with 2 % uranyl acetate. They were then imaged on a JEOL JEM-1400plus TEM (JEOL, Akishima, Tokyo, Japan) at 120 kV of accelerating voltage with a TVIPS f416 camera (TVIPS, Gauting, Germany).

#### 4.4.4. Liposome flotation assay

Lipids used were DOPC (1,2-dioleoyl-sn-glycero-3-phosphocholine), DOPG (1,2-dioleoyl-sn-glycero-3-phospho-(1’-rac-glycerol)), and Rhodamine-PE (1,2-dioleoyl-sn-glycero-3-phosphoethanolamine-N-(lissamine rhodamine B sulfonyl)), all of them purchased from Avanti Polar Lipids (Alabaster, Alabama, USA). The lipids, dissolved in chloroform, were mixed to final ratios (mol:mol) of 69.9 DOPC : 30 DOPG : 0.1 Rhodamine-PE, and evaporated in a glass vial to obtain a thin lipid film. Lipids were resuspended in buffer containing 50 mM HEPES pH 7.5, 50 mM NaCl and 300 mM sucrose, at a final concentration of 5 mg/ml. The lipid film was hydrated for 1 hour and thoroughly vortexed to form multilamellar vesicles. The lipids were then mixed with 0.1 μM of TEV protease and 1 μM of protein of interest. Lipids and protein were left to incubate for 1 hour at RT. The sample was then deposited at the bottom of an ultracentrifuge tube, and mixed with buffer containing sucrose to obtain a bottom layer of 30% of sucrose. Gently, another layer of buffer with 25% of sucrose was deposited, and a final layer of 0% of sucrose on top. In the experiments shown in Figure 1C, the sample was instead applied to the top of the sucrose gradient. Then it was centrifuged at 200,000 g at 4 °C for 30 minutes in a SW 60 Ti Swinging Bucket rotor. All the different fractions of the sucrose gradient were then pipetted out and extra buffer was then added to resuspend the filamented pellet at the bottom. The different fractions were then analysed by SDS PAGE, and the presence of the proteins was visualize either by the fluorescence of the fluorophore-labelled proteins or by staining with Coomassie blue. Experiments with CdvB2ΔC were done with a final concentration of all the proteins of 600nM, and gels were imaged using a GE Amersham™ Typhoon gel imager to image the fluorescent label on the proteins and lipids.

#### 4.4.5. Preparation of lipid in oil suspension for dumbbell-shaped liposome preparations

DOPC, DOPE-PEG2000, DOPG and DOPE-Rhodamine (or DOPE-Atto390 for experiments with proteins with overlapping fluorescence) in chloroform were mixed in a ratio of 93:2:5:0.1 and evaporated in a glass vial under a blow of nitrogen. Lipid mixture was then resolubilized in chloroform to a final concentration of 0.2 mg/ml. A freshly prepared mixture of silicone and mineral oil that was added to the lipids in chloroform slowly dropwise while vortexing gently. After all oil is added to the chloroform, it was vortexed at max speed for 2 minutes and then sonicated for 15 minutes in an ice bath.

#### 4.4.6. Preparation of dumbbell-shaped liposomes with the synthetic membrane shaper

Cdv proteins were mixed in an inner buffer containing 50mM Tris pH 7.5 and 37 % w/v optiprep (Sigma Aldrich, St. Louis, Missouri, USA) to make the solution heavy. In parallel, an outer solution in buffer containing 50 mM Tris pH7.4, 5 mM MgCl2 and glucose to match the osmolarity of the outer solution at 30 mOsm higher than in the inner solution. The DNA nanostars developed in Ref. (24) were then mixed into the outer solution and deposited at the bottom of an imaging chamber. Water in oil droplets of inner buffer containing protein were then formed by pipetting up and down 20 μl of inner solution into 400 μl of oil until a homogeneous droplet size was achieved. The droplets in oil were immediately deposited on top of the outer solution in the imaging chamber, and they were allowed to sediment by gravity through the oil-water interphase. The liposomes were imaged using spinning disk confocal laser microscopy (Olympus IXB1/BX61 microscope, 60× objective, iXon camera) with Andor iQ3 software. Analysis of the images was done with ImageJ (v.2.1.0).

## Acknowledgments

We thank Eli van der Sluis and Ashmiani van den Berg for discussions and protein purification. We acknowledge funding support from the BaSyC program of NWO-OCW and from the ERC Advanced Grant 883684. This research is part of the project no. 2022/45/P/NZ1/01565 co-funded by the National Science Centre and the European Union Framework Programme for Research and Innovation Horizon 2020 under the Marie Skłodowska-Curie grant agreement no. 945339.

## References

Azad K, Guilligay D, Boscheron C, Maity S, De Franceschi N, Sulbaran G, Effantin G, Wang H, Kleman JP, Bassereau P, Schoehn G, Roos WH, Desfosses A, Weissenhorn W. Structural basis of CHMP2A-CHMP3 ESCRT-III polymer assembly and membrane cleavage. Nat Struct Mol Biol. 2023 Jan;30(1):81–90.

Bertin A, de Franceschi N, de la Mora E, Maity S, Alqabandi M, Miguet N, di Cicco A, Roos WH, Mangenot S, Weissenhorn W, Bassereau P. Human ESCRT-III polymers assemble on positively curved membranes and induce helical membrane tube formation. Nat Commun. 2020 May 29;11(1):2663.

Blanch Jover A, De Franceschi N, Fenel D, Weissenhorn W, Dekker C. The archaeal division protein CdvB1 assembles into polymers that are depolymerized by CdvC. FEBS Lett. 2022 Apr;596(7):958–969.

Caillat C, Macheboeuf P, Wu Y, McCarthy AA, Boeri-Erba E, Effantin G, Göttlinger HG, Weissenhorn W, Renesto P. Asymmetric ring structure of Vps4 required for ESCRT-III disassembly. Nat Commun. 2015 Dec 3;6:8781.

Caspi Y, Dekker C. Dividing the Archaeal Way: The Ancient Cdv Cell-Division Machinery. Front Microbiol. 2018 Mar 2;9:174.

Chiaruttini N, Redondo-Morata L, Colom A, Humbert F, Lenz M, Scheuring S, Roux A. Relaxation of Loaded ESCRT-III Spiral Springs Drives Membrane Deformation. Cell. 2015 Nov 5;163(4):866–79.

Cox CJ, Foster PG, Hirt RP, Harris SR, Embley TM. The archaebacterial origin of eukaryotes. Proc Natl Acad Sci U S A. 2008 Dec 23;105(51):20356–61.

De Franceschi N, Alqabandi M, Miguet N, Caillat C, Mangenot S, Weissenhorn W, Bassereau P. The ESCRT protein CHMP2B acts as a diffusion barrier on reconstituted membrane necks. J Cell Sci. 2018 Aug 3;132(4):jcs217968.

De Franceschi N, Barth R, Meindlhumer S, Fragasso A, Dekker C. Dynamin A as a one-component division machinery for synthetic cells. Nat Nanotechnol. 2024 Jan;19(1):70–76.

De Franceschi N, Pezeshkian W, Fragasso A, Bruininks BMH, Tsai S, Marrink SJ, Dekker C. Synthetic Membrane Shaper for Controlled Liposome Deformation. ACS Nano. 2022 Nov 28;17(2):966–78.

Dobro MJ, Samson RY, Yu Z, McCullough J, Ding HJ, Chong PL, Bell SD, Jensen GJ. Electron cryotomography of ESCRT assemblies and dividing Sulfolobus cells suggests that spiraling filaments are involved in membrane scission. Mol Biol Cell. 2013 Aug;24(15):2319–27.

Harker-Kirschneck L, Hafner AE, Yao T, Vanhille-Campos C, Jiang X, Pulschen A, Hurtig F, Hryniuk D, Culley S, Henriques R, Baum B, Šarić A. Physical mechanisms of ESCRT-III-driven cell division. Proc Natl Acad Sci U S A. 2022 Jan 4;119(1):e2107763119.

Hurley JH. ESCRTs are everywhere. EMBO J. 2015 Oct 1;34(19):2398–407.

Hurtig F, Burgers TCQ, Cezanne A, Jiang X, Mol FN, Traparić J, Pulschen AA, Nierhaus T, Tarrason-Risa G, Harker-Kirschneck L, Löwe J, Šarić A, Vlijm R, Baum B. The patterned assembly and stepwise Vps4-mediated disassembly of composite ESCRT-III polymers drives archaeal cell division. Sci Adv. 2023 Mar 17;9(11):eade5224.

Lata S, Schoehn G, Jain A, Pires R, Piehler J, Gottlinger HG, Weissenhorn W. Helical structures of ESCRT-III are disassembled by VPS4. Science. 2008 Sep 5;321(5894):1354–7.

Lee IH, Kai H, Carlson LA, Groves JT, Hurley JH. Negative membrane curvature catalyzes nucleation of endosomal sorting complex required for transport (ESCRT)-III assembly. Proc Natl Acad Sci U S A. 2015 Dec 29;112(52):15892–7.

Lindås AC, Karlsson EA, Lindgren MT, Ettema TJ, Bernander R. A unique cell division machinery in the Archaea. Proc Natl Acad Sci U S A. 2008 Dec 2;105(48):18942–6.

Makarova KS, Yutin N, Bell SD, Koonin EV. Evolution of diverse cell division and vesicle formation systems in Archaea. Nat Rev Microbiol. 2010 Oct;8(10):731–41.

Moriscot C, Gribaldo S, Jault JM, Krupovic M, Arnaud J, Jamin M, Schoehn G, Forterre P, Weissenhorn W, Renesto P. Crenarchaeal CdvA forms double-helical filaments containing DNA and interacts with ESCRT-III-like CdvB. PLoS One. 2011;6(7):e21921.

Pfitzner AK, Mercier V, Jiang X, Moser von Filseck J, Baum B, Šarić A, Roux A. An ESCRT-III Polymerization Sequence Drives Membrane Deformation and Fission. Cell. 2020 Sep 3;182(5):1140–1155.e18.

Pulschen AA, Mutavchiev DR, Culley S, Sebastian KN, Roubinet J, Roubinet M, Risa GT, van Wolferen M, Roubinet C, Schmidt U, Dey G, Albers SV, Henriques R, Baum B. Live Imaging of a Hyperthermophilic Archaeon Reveals Distinct Roles for Two ESCRT-III Homologs in Ensuring a Robust and Symmetric Division. Curr Biol. 2020 Jul 20;30(14):2852–2859.e4.

Samson RY, Obita T, Freund SM, Williams RL, Bell SD. A role for the ESCRT system in cell division in archaea. Science. 2008 Dec 12;322(5908):1710–3.

Samson RY, Obita T, Hodgson B, Shaw MK, Chong PL, Williams RL, Bell SD. Molecular and structural basis of ESCRT-III recruitment to membranes during archaeal cell division. Mol Cell. 2011 Jan 21;41(2):186–96.

Shim S, Kimpler LA, Hanson PI. Structure/function analysis of four core ESCRT-III proteins reveals common regulatory role for extreme C-terminal domain. Traffic. 2007 Aug;8(8):1068–79.

Schöneberg J, Lee IH, Iwasa JH, Hurley JH. Reverse-topology membrane scission by the ESCRT proteins. Nat Rev Mol Cell Biol. 2017 Jan;18(1):5–17.

Tang S, Henne WM, Borbat PP, Buchkovich NJ, Freed JH, Mao Y, Fromme JC, Emr SD. Structural basis for activation, assembly and membrane binding of ESCRT-III Snf7 filaments. Elife. 2015 Dec 15;4:e12548.

Tarrason Risa G, Hurtig F, Bray S, Hafner AE, Harker-Kirschneck L, Faull P, Davis C, Papatziamou D, Mutavchiev DR, Fan C, Meneguello L, Arashiro Pulschen A, Dey G, Culley S, Kilkenny M, Souza DP, Pellegrini L, de Bruin RAM, Henriques R, Snijders AP, Šarić A, Lindås AC, Robinson NP, Baum B. The proteasome controls ESCRT-III-mediated cell division in an archaeon. Science. 2020 Aug 7;369(6504):eaaz2532.

