## Supplementary Figures 1,2,3 for "Interaction hierarchy among Cdv proteins drives recruitment to membrane necks"

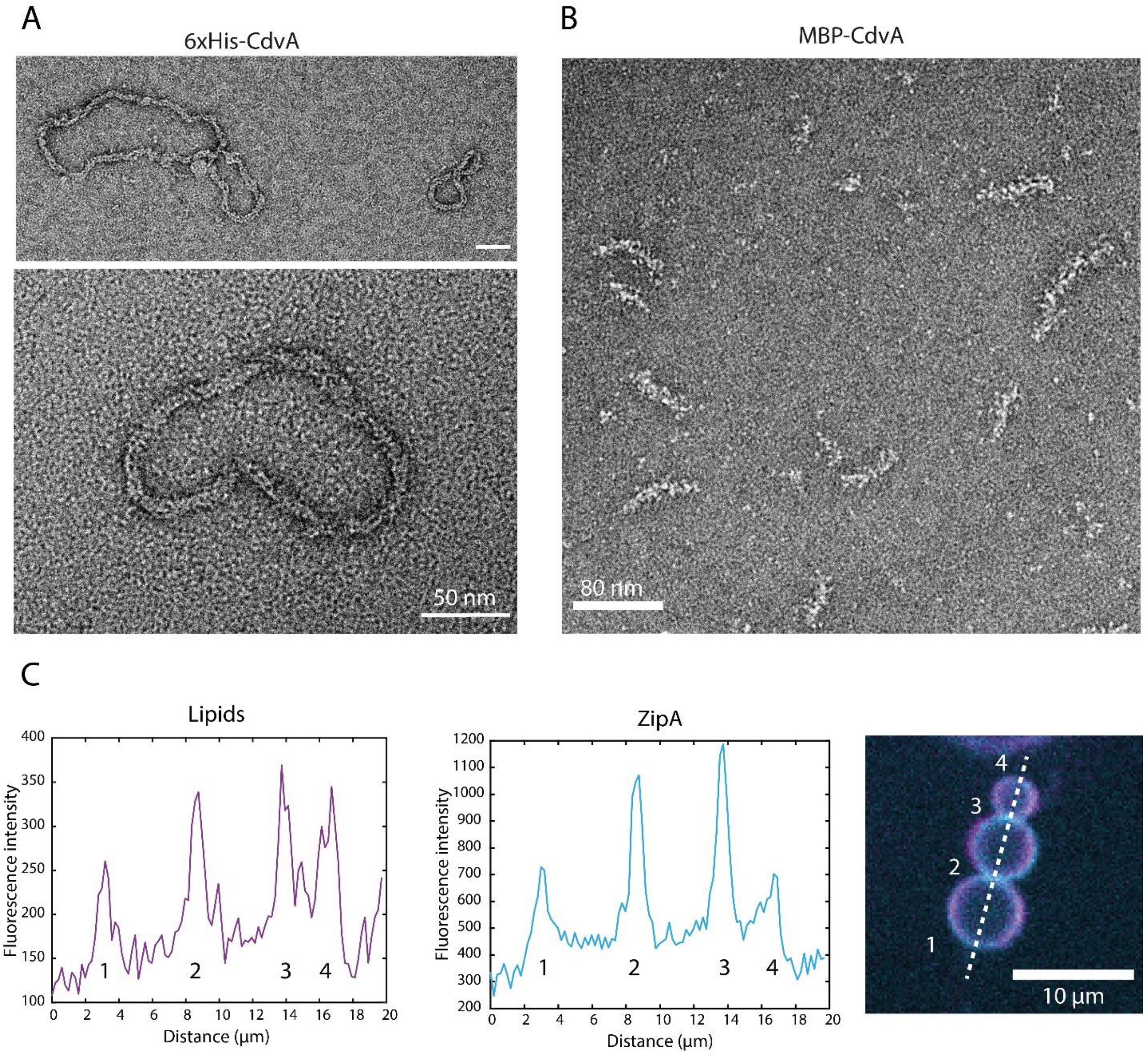


**Supplementary Figure 1: (A)** CdvA purified through His-Tag purification as described in Moriscot et al. 2011. **(B)** Short and thick polymers formed by polymerization of MBP-CdvA. **(C)** line scan profile of ZipA and fluorescent lipids, showing similar patterns and absence of protein enrichment at necks. Scale Bar: 10 µm.

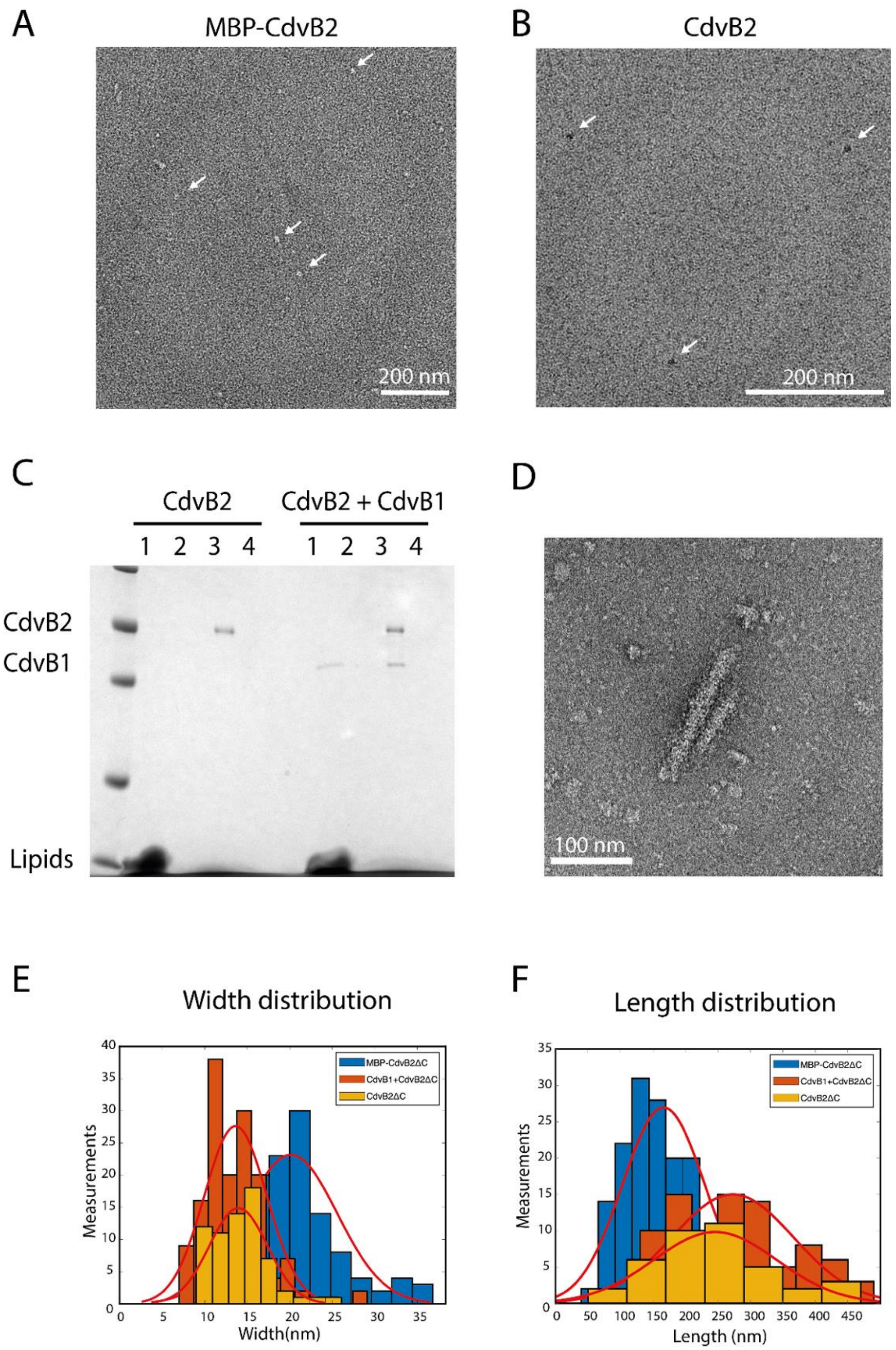


**Supplementary Figure 2: (A)** Absence of filament formation by MBP-CdvB. **(B)** Absence of filament formation by CdvB after MBP removal. **(C)** Absence of membrane binding by CdvB2 full-length, neither alone nor together with CdvB1. **(D)** Negative staining TEM images of MBP-CdvB2ΔC, showing short linear filaments. **(E,F)** Width and length distribution of the polymers formed by MBP- CdvB2ΔC, CdvB2ΔC and CdvB1 + CdvB2ΔC.

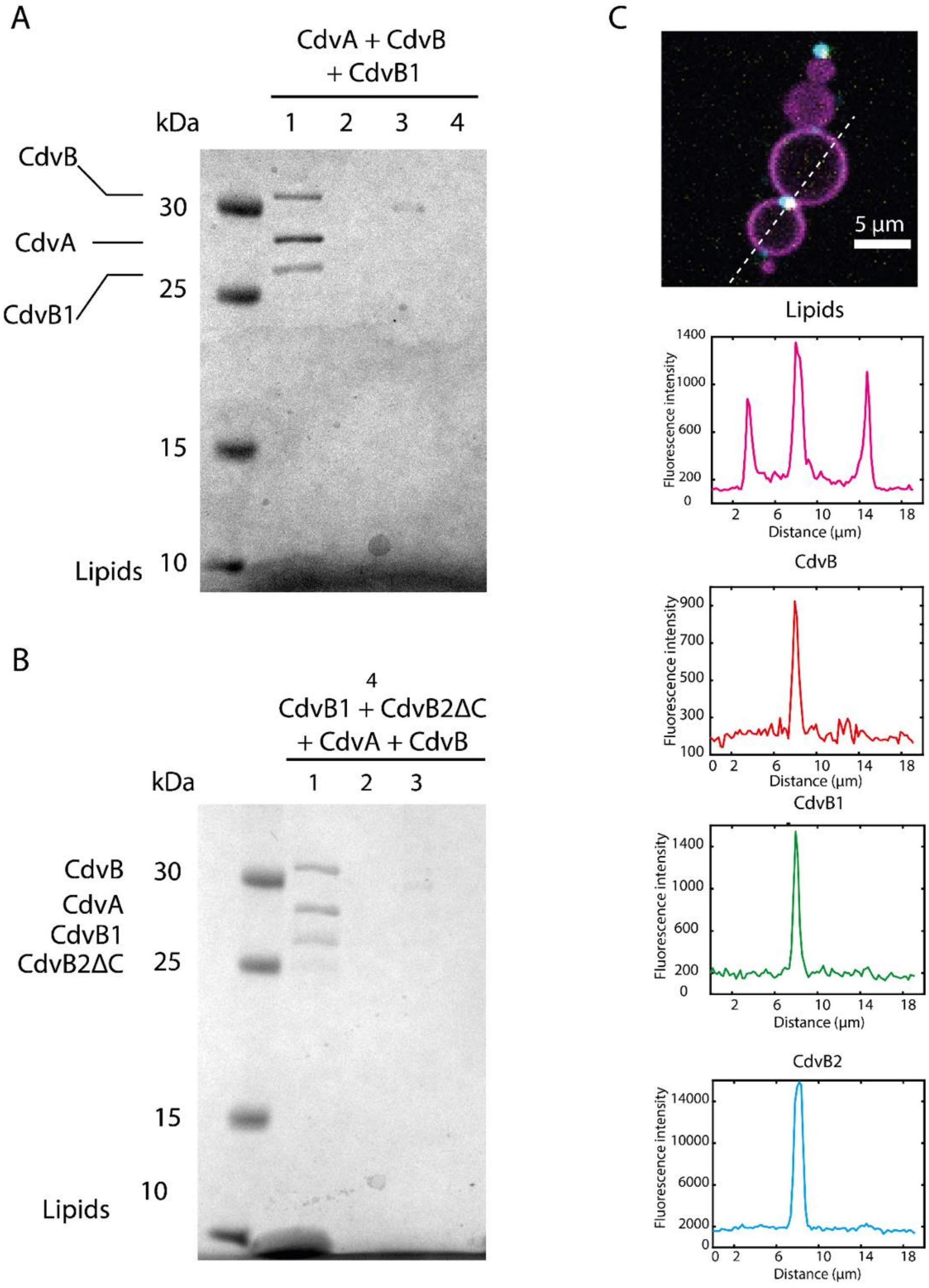


**Supplementary Figure 3: (A)** Coomassie staining showing co-recruitment of CdvA, CdvB and CdvB1 to liposomes. **(B)** Coomassie staining showing co-recruitment of CdvA, CdvB, CdvB1 and CdvB2ΔC to liposomes. **(C)** Fluorescence intesity profile across the dotted line of a quaternaty complex including CdvA, CdvB, CdvB1 and CdvB2ΔC reconstituted in dumbbell liposomes, showing clear enrichment of all components at the neck.
